# Infectious myonecrosis virus (IMNV) and decapod iridescent virus 1 (DIV1) detected in *Penaeus monodon* from the Indian Ocean

**DOI:** 10.1101/2020.09.19.304972

**Authors:** Jiraporn Srisala, Piyachat Sanguanrut, Saensook Laiphrom, Jittima Siriwattano, Juthatip Khudet, Dararat Thaiue, Sorawit Powtongsook, Timothy W. Flegel, Kallaya Sritunyalucksana

## Abstract

Infectious myonecrosis virus (IMNV) was first discovered in the Americas in 2004 as a new lethal pathogen of cultivated whiteleg shrimp *Penaeus vannamei*, but infections were not lethal for the giant tiger shrimp *Penaeus monodon*. In 2007, it was reported in diseased *P. vannamei* cultivated in Indonesia but, until recently, not from other countries in Asia. Decapod iridescent virus (DIV1) was first reported from China in 2016 and is lethal for the crayfish *Cherax quadricarinatus* and *Procambarus clarkii*, for the penaeid shrimp *P. vannamei* and *P. chinensis* and for the palaemonid shrimp *Macrobrachium rosenbergii* and *Exopalaemon carinicauda*. It has not yet been reported from other Asian countries. Here we describe the occurrence of positive test results for IMNV and DIV1 using polymerase chain reaction (PCR) technology during screening of grossly normal, broodstock-size, wild *P. monodon* captured from the Indian Ocean and held in a biosecurity facility for screening. Amplicons for each virus were obtained from two widely separated targets in the relevant viral genomes listed at GenBank, and sequencing revealed 99-100% identity to the targets for each virus. Based on these results, the captured specimens were destroyed. The results raised the possibility that grossly normal, captured *P. monodon* might serve as potential vehicles for introduction of IMNV and/or DIV1 to shrimp hatcheries and farms. Thus, we recommend that appropriate precautions be taken to avoid this possibility.

## INTRODUCTION

Infectious myonecrosis virus (IMNV) was first described from the Americas as a lethal pathogen of the whiteleg shrimp *Penaeus vannamei* (Poulos Lightner, 2006; Poulos, et al., 2006). However, it was also infectious but not lethal for *P. stylirostris* and *P. monodon* (Poulos Lightner, 2006; Poulos, et al., 2006; Tang, et al., 2005). It was subsequently introduced to Indonesia around 2006 (Senapin, et al., 2007) but has been slow to spread to other Asian countries (Sahul Hameed, et al., 2017; Senapin, et al., 2011). Decapod iridescent virus (DIV1)(Chen, et al., 2019) in *Exopalaemon carinicauda* was first described from China as *Cherax quadricarinatus* iridovirus/CQIV infectious for *C. quadricarinatus, Procambarus clarkii* and *P. vannamei* (Li, et al., 2017; Xu, et al., 2016) or as shrimp hemocyte iridescent virus (SHIV)(Qiu, et al., 2017; Qiu, et al., 2018) infectious for *P. vannamei, P. chinensis* and *Macrobrachium rosenbergii*. Thus, DIV1 has a wide known-host range that includes several economically important cultured species.

Infectious myonecrosis disease caused by IMNV results in gross signs of disease characterized by whitening of the skeletal muscles, especially in the abdominal region of affected shrimp (Poulos, et al., 2006). When such tissues are examined for histopathology, muscle lesions can be seen that are characterized by myonecrosis, hemocyte aggregation and the presence of basophilic, cytoplasmic inclusions (Poulos & Lightner, 2006; Poulos, et al., 2006). There are also published nested, reverse-transcriptase, polymerase chain reaction (RT-PCR) methods for its detection, even in lightly-infected specimens that may show no gross signs or histological signs of infection (Poulos & Lightner, 2006; Senapin, et al., 2007). The presence of IMNV nucleic acid (RNA) in the cytoplasm of cells in muscle lesions has been confirmed by *in situ* hybridization (ISH) assays (Poulos, et al., 2006).

Disease caused by DIV1 results in massive mortality accompanied by gross signs of disease characterized by features including an empty stomach and midgut, a pale hepatopancreas and a soft shell (Qiu, et al., 2017). Histological signs of DIV1 include pathognomonic lesions characterized by the presence of unique, lightly basophilic viral inclusions in the cytoplasm of cells of the hematopoietic tissue (HPT) (Qiu, et al., 2017). The same specimens also show severe necrosis of the lymphoid organ (LO) characterized by loss of tubule structure and the presence of basophilic, cytoplasmic inclusions including karyorrhetic and pyknotic nuclei (Sanguanrut et al., 2020). However, in lightly-infected shrimp, DIV1 may also be detected using nested PCR. The presence of DIV1-DNA in the cytoplasm of cells in these tissues has been confirmed by ISH (Qiu, et al., 2017). It also reveals presence of the virus in the cytoplasm of cells of the antennal gland and of connective tissues including those of the skeletal muscles, the subcuticulum, the hepatopancreas (HP) and the anterior midgut cecum (AMC).

IMNV has posed a threat to Asian aquaculture since its introduction to Indonesia around 2006 but it had not spread from there (Senapin, et al., 2011) until relatively recently (Sahul Hameed, et al., 2017). Thus, it remains a threat to other countries from which it has not yet been reported. The recent description of DIV1 from China and its pathogenicity for several crustacean species also poses a new threat to all other shrimp culturing countries from which it has not yet been reported.

In a program to establish a breeding stock from wild, captured specimens of *P. monodon* from the Indian Ocean, we participated by screening individuals from captured batches for a list of 14 known viral pathogens and parasites by non-destructive polymerase chain reaction (PCR) methods while they were being held in a biosecure facility. It is important to understand that this was not an epidemiological survey. Thus, there was no geographical collection plan and shrimp were sequentially tested for each pathogen such that a positive test for any specimen at any stage of testing resulted in its destruction and in no on-going testing of its nucleic acid extracts for pathogens for which it had not yet been tested. In other words, not every shrimp collected was subjected to screening for every pathogen in the screening list. As a result, the information gained from this study cannot be used to estimate the possible prevalence of IMNV and DIV1 positive shrimp in wild populations of *P. monodon* in the Indian Ocean.

We detected IMNV and DIV1 in both batches captured in April 2018 and March 2019. Here we describe details of the methods used and results obtained. Although we carried out no bioassays using the positive specimens as a source of inoculum, our results open the possibility that grossly normal, captured *P. monodon* might serve as potential vehicles for introduction of IMNV and/or DIV1 to shrimp hatcheries and farms. Thus, we recommend that appropriate precautions be taken to avoid this possibility.

## MATERIALS AND METHODS

### Shrimp specimens and sampling

Thai fishermen were hired to capture broodstock size specimens of the giant tiger shrimp P. monodon from the Indian Ocean. These were transported to a biosecure facility where they were held individually to be screened for a list of 14 viral pathogens using PCR technology. Two lots of shrimp (Lot 1 of 14 shrimp and Lot 4 of 76 shrimp) are relevant to this study. In the total of 90 shrimp analyzed. These lots were received in April 2018 and March 2019. The shrimp were held individually in foam boxes (approximately 35 x48×35 supplied with recirculating seawater at 30-32 ppt and 20-22oC. They were fed with a commercial shrimp feed at 1-2% per gram shrimp, 10 times per day, and excess, uneaten feed was removed once a day.

The experimental protocol was approved by the Shrimp Genetic Improvement Center (SGIC), National Center for Genetic Engineering and Biotechnology (BIOTEC) and National Science and Technology Development Agency (NSTDA).

Nucleic acid templates to be used in PCR testing for IMNV and DIV1 were extracted from the tips of pleopods (abdominal swimming appendages). For pleopod clipping, approximately 20 mg of tissue was clipped and homogenized in DNA lysis buffer and trizol reagent and was transferred to the laboratory for nucleic acid extraction within 3 days. Specimens from Lot 1 were tested by PCR methods only. However, shrimp positive for IMNV and DIV1 from Lot 4 were stunned in ice water and fixed with Davidson’s fixative for standard histological analysis using hematoxylin and eosin (H&E) staining (Bell&Lightner, 1988) and for *in situ* hybridization (ISH) analysis.

### Preparation of DNA and RNA templates

For DNA extraction, pleopod specimens collected from each shrimp were individually homogenized in DNA lysis buffer (50 mM Tris-base, 100 mM EDTA, 50 mM NaCl, 2% (w/v) SDS and 100 µg/ml proteinase K) and incubated at 56°C for 1 h before extraction using a QIAamp DNA Mini Kit (Qiagen, Germany) according to the manufacturer’s directions. For RNA extraction, pleopods were homogenized in 1 ml of Trizol Reagent (Invitrogen, USA) and extracted following the Trizol reagent protocol. The RNA pellet was resuspended with 30 µl of DNase/RNase free water and digested with DNase I (NEB) following the manufacturer protocol. Subsequently, the sample was re-extracted by the same method. Total DNA and RNA concentration were determined by Qubit 3.0 Fluorometer (Life Technologies)

### RT-PCR methods for IMNV

Two methods were used for RT-PCR detection of IMNV. One followed the original protocol (Poulos Lightner, 2006; Poulos, et al., 2006) that is also the recommended method of the World Organization for Animal Health (OIE) in its Manual of Diagnostic Tests for Aquatic Animals (Anonymous, 2017). It targets the major capsid protein (MCP) gene of IMNV. Here it is called the IMNV-O method. The other followed a later publication (Senapin, et al., 2007) (here called the IMNV-S method). It targets the RNA-dependent, RNA polymerase (RdRp) gene of IMNV. Briefly for the IMNV-O method, the first step primers were 4587F: 5’-CGA CGCTGCTAACCATACAA-3’ and 4914R: 5’-ACTCGGCTGTTCGATCAAGT-3’. The reaction mix contained 1X Reaction Mix (Invitrogen), 0.4 µM each primer, 1 µl of SuperScript III RT/Platinum Taq Mix (Invitrogen, USA) and 20 ng of RNA template in 25 µl total reaction volume. The PCR protocol was 50°C for 30 min, 94°C for 2 min, followed by 35 cycles of 94°C for 30 sec, 60°C for 30 sec and 68°C for 45 sec followed by extension at 68°C for 5 min. The nested step primers were 4725NF:5’-GGCACATGCTCAGAGACA3’ and 4863NR: 5’-AGCGCTGAGTCCAGTCT TG-3’ and the reaction mix contained 1 μl of first PCR product, 1X OneTaq Hot Start Master Mix (NEB), 0.2 µM each primer in a total volume of 25 µl. The PCR protocol was 94°C for 5 min, followed by 30 cycles of 94°C for 30 sec 60°C for 30 sec and 72°C for 30 sec followed by extension at 72°C for 5 min. The amplicons yielded were 328 bp and 139 bp, respectively.

For IMNV-S, the first step primers were IMNV F13: 5’-TTTATACACCGCAAGAATTGG CCAA-3’ and IMNV R13: 5’-AGATTTGGGAGATTGGGTCGTATCC-3’ with an expected amplicon of 600 bp. The nested step primers were IMNV F13N: 5’-TGTTTATGCTTGGGA TG GAA-3’ and IMNV R13N: 5’-TCGAAAGTTGTTGGCTGATG-3’ with an expected amplicon of 282 bp. First step reaction mix contained 1X Reaction Mix (Invitrogen), 0.4 µM each primer, 1 µl of SuperScript III RT/Platinum Taq Mix (Invitrogen, USA) and 20 ng of RNA template in a total reaction volume of 25 µl. The PCR protocol was 50°C for 30 min, 94°C for 2 min, followed by 35 cycles of 94°C for 30 sec, 45°C for 30 sec and 72°C for 45 sec followed by extension at 72°C for 5 min. The nested PCR mix contained 1 μl of first PCR product, 1X OneTaq Hot Start Master Mix (NEB) and 0.2 µM each primer in a total volume of 25 µl. The PCR protocol was 94°C for 5 min, followed by 30 cycles of 94°C for 30 s, 60°C for 30 sec and 68°C for 30 sec followed by extension at 68°C for 5 min. When RT-PCR amplicons were detected by agarose gel electrophoresis, they were cut from gels, purified, cloned and sent for sequencing by Macrogen, Korea. The sequences were then analyzed by the tools available at National Center for Biotechnology Information (NCBI).

### PCR methods for DIV1

Two primer sets were used for DIV1 detection; ATPase and MCP primer set at the different genome target regions. The ATPase primers were followed those described by Qiu et al. 2017 with some modification. The semi-nested PCR profile with the primers SHIV-F1 and SHIV-R1 for the first step PCR with the expected amplicon of 457 bp and the primers SHIV-F1 and SHIV-R2 for the second step PCR with the expected amplicon of 213 bp. The second set was designed from the GenBank record of the DIV1 major capsid protein (MCP) as an in-house, confirmatory method. The primers for the first step reaction were DIV1-F576: 5’-TAGCAGCTTCGGAGCATTGA-3’ and DIV1-R576: 5’-GCAAGGTTCCTCAGG TTGGA-3’ with an expected amplicon size of 576 bp. The primers for nested step were DIV1-F409: 5’-TAATCGGCAGTCATCACGGG-3’ and DIV1-R576: 5’-GCAAGGTTCCTCAGG TTGGA-3’ with an expected amplicon size of 409 bp. The first step PCR reaction mixture contained 1 μl of DNA template, 1X OneTaq Hot Start Master Mix (NEB), 0.4 µM each primer in total volume of 25 µl. The PCR protocol was 94°C for 5 min, followed by 35 cycles of 94°C for 30 sec, 58°C for 30 sec and 72°C for 45 sec followed by extension at 72°C for 5 min. The semi-nested reaction mixture contained 1 μl of first PCR product, 1X OneTaq Hot Start Master Mix (NEB) and 0.2 µM each primer in a total volume of 25 µl. The PCR protocol was at 94°C for 5 min, followed by 25 cycles of 94°C for 30 sec, 58°C for 30 sec and 72°C for 30 sec followed by extension at 72°C for 5 min.

### Amplicon sequencing and analysis

PCR amplicons were cloned and sent to Macrogen, Korea for sequencing. The sequences were analyzed by tools available at the website of the National Center for Biotechnology Information (NCBI). For each sample sent for sequencing, at least 3 clones were sequenced from both strands and a consensus sequence was obtained via multiple alignment of the results for each sample. When a difference occurred at 1 position in one strand of the alignment but was the same in two or more remaining strands, the latter base was included in the consensus sequence. This happened not more than 3 times for any consensus sequence.

### Histological analysis and *in situ* hybridization (ISH) assays

Living shrimp were stunned in a seawater ice bath before being injected with Davidson’s fixative and processed for embedding in paraffin blocks in order to cut tissue section for staining with hematoxylin and eosin (H&E) as previously described (Bell Lightner, 1988). Only specimens from Lot 4 that gave positive test results for either IMNV or DIV1 were processed further for histological analysis. *In situ* hybridization assays were carried out using adjacent tissue sections from the same paraffin blocks used for H&E-stained sections. For both IMNV and DIV1, H&E-stained tissue sections from the respective shrimp specimens were examined for the presence of the characteristic lesions for each pathogen as described in the introduction section. Positive control material for ISH assays for DIV1 consisted of microscope slides prepared from paraffin blocks derived from a laboratory challenge test with *P. vannamei*. The positive control material for ISH assays for IMNV consisted of a paraffin block of IMNV-infected tissue of *P. vannamei* purchased from the University of Arizona. Aquaculture Pathology Laboratory.

### For DIV1-ISH testing

The PCR primer pairs of the first step of the ATPase gene detection method (amplicon size 457 bp) and the MCP gene detection method (amplicon size 576 bp) were used to prepare DIG-labeled DIV1 probes. Briefly, The DNA probes were generated using a PCR DIG labeling kit (Roach, Germany) followed by purification using a Gel/PCR cleanup kit (Geneaid, Taiwan) according to the manufacturer’s directions. The ISH protocol was performed as follows: the slides of adjacent tissue sections were incubated at 60°C for 1 h, deparaffinized, rehydrated in a series of graded ethanol, distilled water and TNE buffer. Tissue slides were treated with 5µg/ml proteinase K in TNE buffer (500 mM Tris-Cl, 100 mM NaCl, l0 mM EDTA) at 37°C for 15 min in humidified chamber. The slides were incubated with 0.5M EDTA for 1 h, cold 4% formaldehyde for 5 min and distilled water at room temperature (RT) for 5 min. The sections were incubated in pre-hybridized buffer (4X SSC containing 50% (v/v) deionized formamide) at 37°C for at least 10 min. Then, 100-200 ng of each DIG-labeled probe, was mixed with hybridization buffer (50% deionized formamide, 5% (w/v) dextran sulfate, 1X Denhardt’s solution (Sigma, USA), 0.25 mg/ml salmon sperm DNA (Invitrogen) and 4X SSC before denaturation at 95°C for 5 min followed by immediate chilling on ice. The denatured probes were then pipetted onto each slide and covered with a cover slip after which the slides were heated at 95°C for 7 min. They were then incubated at 42°C overnight in a humidified chamber. The slides were washed in 2X SSC at 37°C, 1X SSC at 42°C, 0.5X SSC at 42°C and then Buffer I (1M Tris– HCL, 1.5M NaCl, pH 7.5) for 5 min. Next, the slides were incubated with 0.5% blocking buffer (Roche, Germany) in 1X Buffer I at RT for 1 h before Incubation with 1:500 anti-DIG-AP antibody solution at 37°C for 1 h and final washing with 2×10 min Buffer I. The signal was developed using NBT/BCIP solution (Roche, Germany) in a dark chamber at RT after which each slide was counterstained with 0.5% Bismarck Brown Y (Sigma, USA) before dehydration, addition of a drop of Permount (Fisher Scientific, USA) and covering with a cover glass for examination by light microscopy.

## RESULTS

### Order of presentation and overview of PCR and histological results

From shrimp Lot 1 (14 specimens, April 2018), 2 specimens gave positive RT-PCR test results for IMNV and 5 gave positive test results for DIV1. No samples were prepared for histological analysis for specimens of Lot 1. From shrimp Lot. 4 (76 specimens, March 2019), 4 samples were RT-PCR positive for IMNV and 8 were PCR positive for DIV1. None of the specimens were positive for both IMNV and DIV1. In the following sections, results from the shrimp positive for IMNV (Lots 1 and 4) will be covered first, followed by results for DIV1 from Lot 4. No sequencing was done with the IMNV positive samples in Lot 4 because they were positive by one PCR method only. However, histological results and ISH test results are presented. For DIV1 samples, PCR results, amplicon sequencing results, histology results and ISH results are presented.

### Positive RT-PCR results and amplicon sequencing for IMNV from Lot 1

In specimen Lot 1, using the IMNV-O method for IMNV detection, we obtained positive nested RT-PCR test results for 2 specimens from 26 tested (**Fig. 1**). The amplicons were sequenced and aligned with the homologous region of the MCP gene from 3 full IMNV reference sequences at GenBank (**Fig. 2**). One reference sequence was from the first report of IMNV in Brazil in 2006 (GenBank AY570982.3) (Poulos, et al., 2006), another from Brazil in 2014 (KJ556923.1) and yet another from Indonesia in 2007 (EF061744.1). The alignment revealed that our sequences from specimens 1 & 2 were identical to one another, as were the two sequences from Brazil. Excluding the primer sequences outlined in Fig. 2, our two sequences from *P. monodon* differed from the two Brazilian sequences for only 1 base in 288, giving a sequence identity of 287/288 = 99.7%. In contrast, they differed by 2 bases with the sequence from Indonesia, giving a sequence identity of 99.3%. In contrast, the deduced amino acid sequences for all five records were 100% identical, indicating that the base differences represented synonymous mutations.

**Figure 1.**
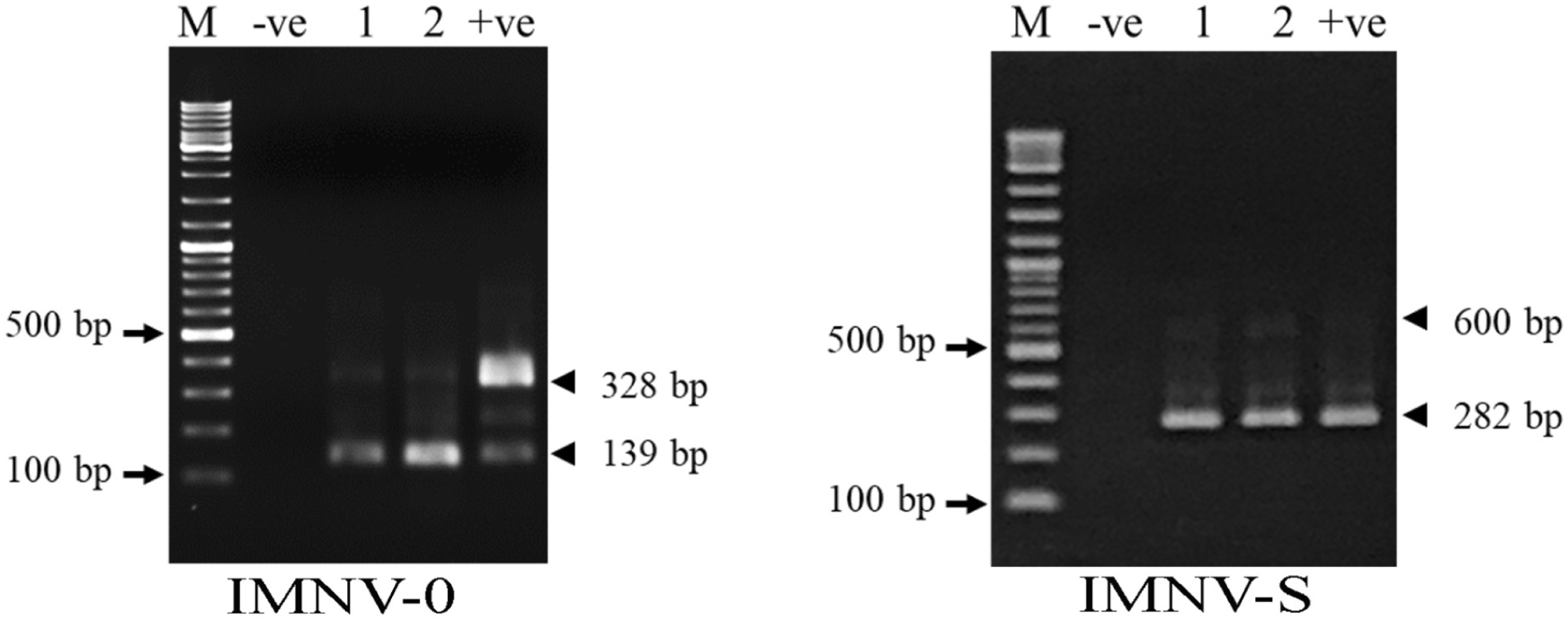
Photographs of agarose gels to detect amplicons from use of the IMNV-O and IMNV-S methods for 2 shrimp specimens (1 & 2) from Lot 1. Both specimens were positive with both methods. M = molecular marker; -ve = negative control; +ve = positive control.

**Figure 2.**
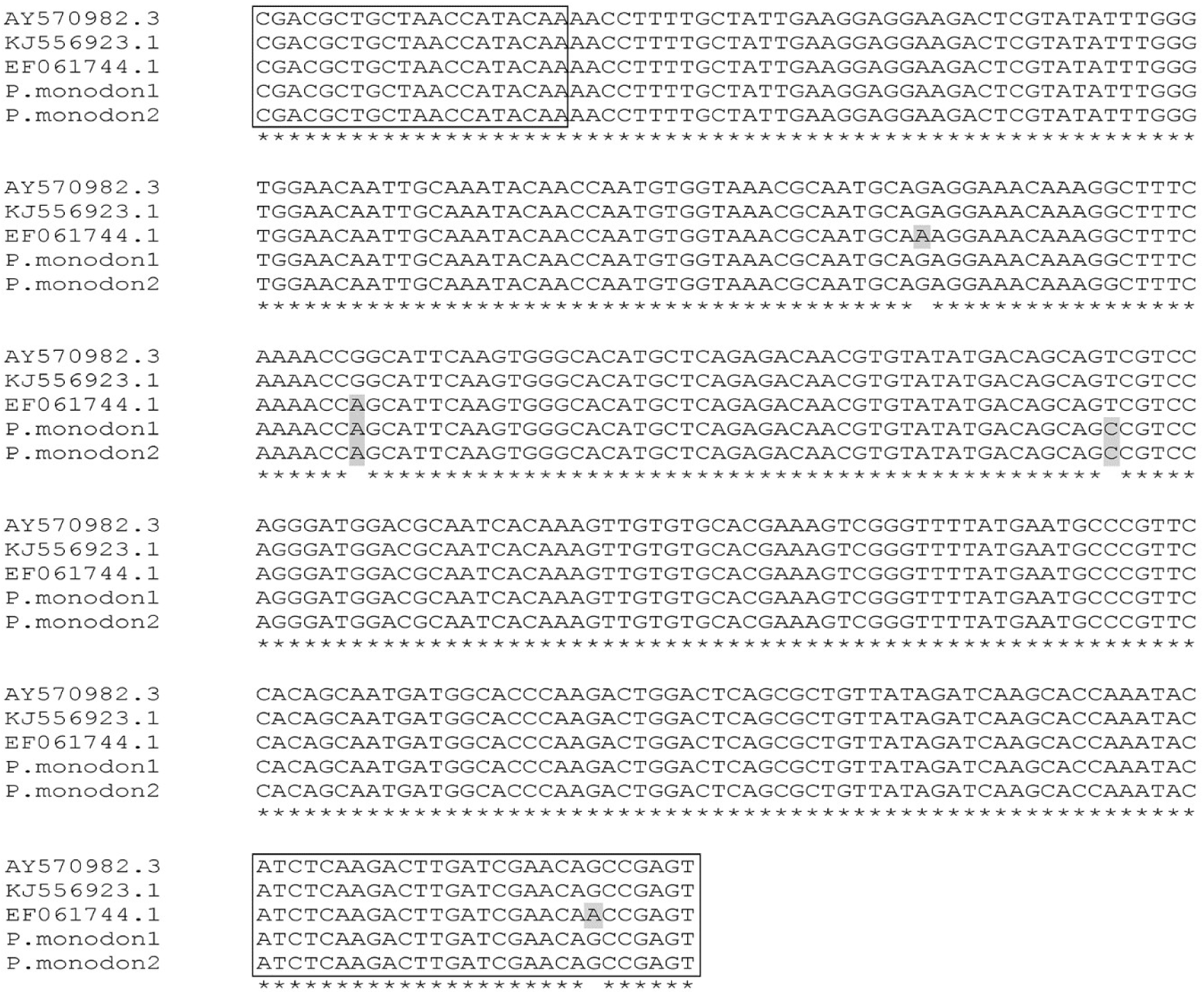
Multiple alignment of the IMNV-O amplicon sequences from our *P. monodon* specimens 1 & 2 in shrimp Lot 1 with the matching region of the MCP gene of IMNV in GenBank records AY570982.3 (the first IMNV sequence from Brazil in 2006), EF061744.1 (from Indonesia in 2007) and KJ556923.1 (from Brazil in 2014). Bases in grey outline indicate differences from the original sequence of AY570982.3 (top row in the alignment). The areas in boxes indicate the primer sequences. Also shown is and alignment of the deduced amino acid sequences showing 100% identity for all.

Using the IMNV-S method for IMNV detection with the 2 specimens in Lot 1, we obtained positive RT-PCR results (Fig. 1) for the same 2 specimens that gave positive results with the IMNV-O method above. The 2 amplicons were sequenced and aligned with the homologous regions of the RdRp gene from the same GenBank reference sequences as used above for the MCP gene. The results revealed that the sequences from our *P. monodon* specimens 1 & 2 were identical to one another but differed from the original GenBank sequence by 5 bases, resulting in 237/242 = 97.9% identity (i.e., excluding the primer sequences) (**Fig. 3**). However, translation to the deduced amino acid sequences followed by alignment revealed that 4 of the differences constituted synonymous mutations in the RdRp gene. The one exception was the change from glutamic acid (E) to lysine (K), denoted by Clustal 2.1 as a conservative replacement. We found it curious that the MCP gene from our samples was more conserved than the RdRp gene, since changes in the latter would seem to be more critical for viral survival than changes in the MCP gene. It is unknown whether this change would affect the virulence of IMNV.

**Figure 3.**
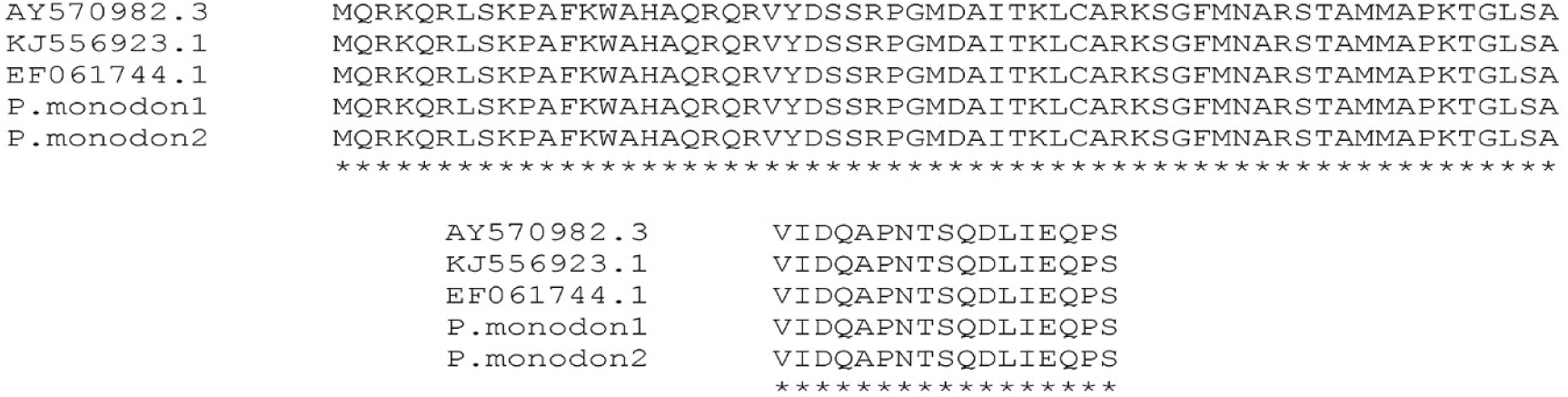
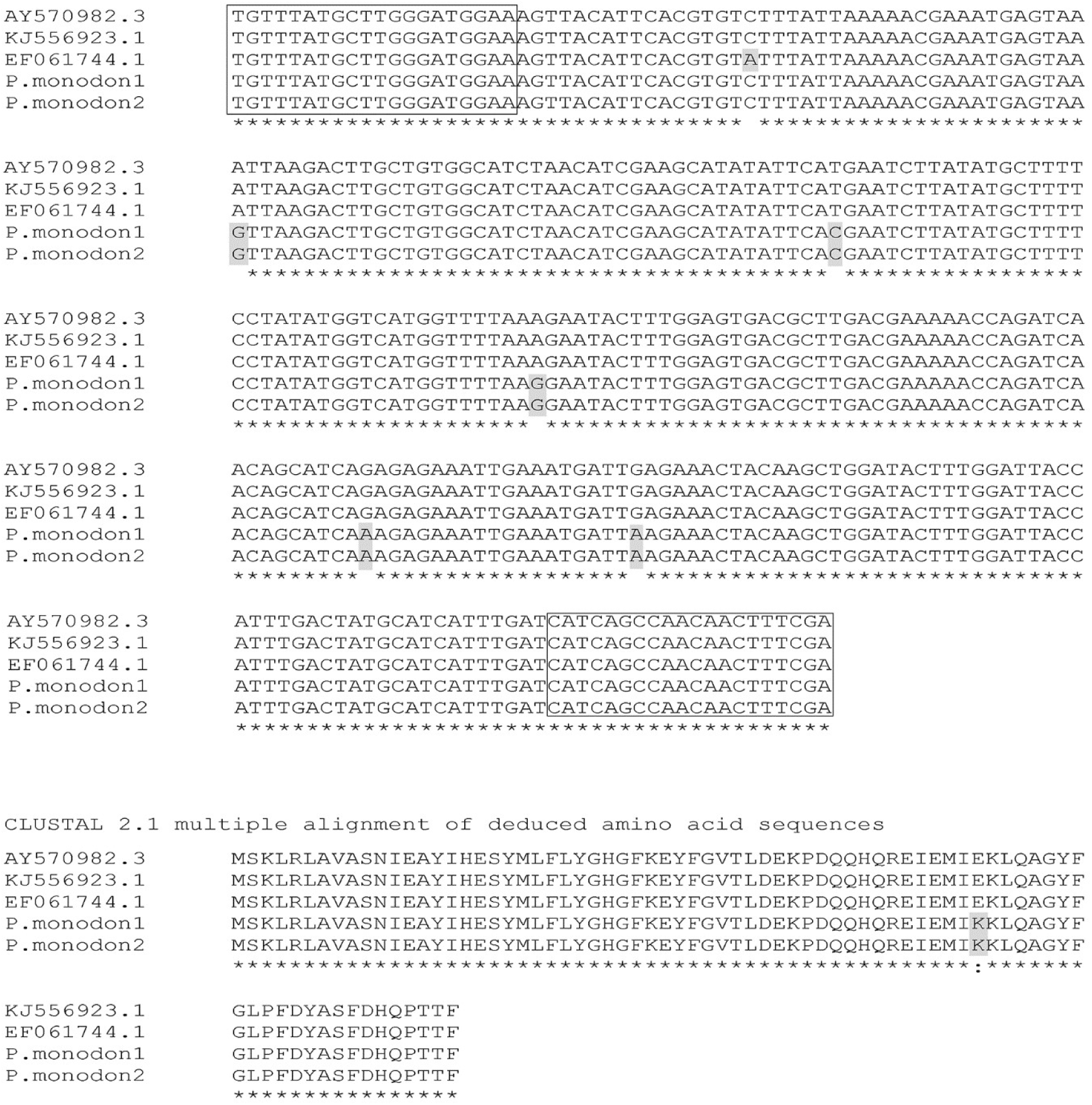
Multiple alignment of the IMNV-S amplicon sequences from specimens 1 & 2 from shrimp Lot 1 with the RdRp gene of IMNV with the same GenBank records used in Fig. 2 above. Bases in white text and black outline indicate difference from the sequence of AY570982.3. Also included is an alignment of the deduced amino acid sequences with only one amino acid difference in the *P. Monodon* samples from the Indian Ocean.

In summary, the 2 IMNV-positive specimens from Lot 1 gave identical amplicons for two different IMNV genes that shared 98-99% identity to the matching sequences from 3 full genome sequences for IMNV isolates from diseased shrimp from Brazil and Indonesia. These results suggest that the 2 *P. monodon* specimens captured from the Indian Ocean wild fishery may have carried an infectious form of IMNV. However, confirmation of this possibility would require at least full genome sequencing and at best bioassays. As stated in the first section of the results, we were unable to do histological or ISH analysis with these specimens.

### Positive RT-PCR results and amplicon sequencing for IMNV from Lot 4

In specimen Lot 4, using the IMNV-O method for IMNV detection, we obtained positive nested RT-PCR test for 4 specimens from 76 tested (**Fig. 4**). In contrast to Lot 1, these 4 specimens gave no amplicons using the IMNV-S method (**Fig. 4**).

**Figure 4.**
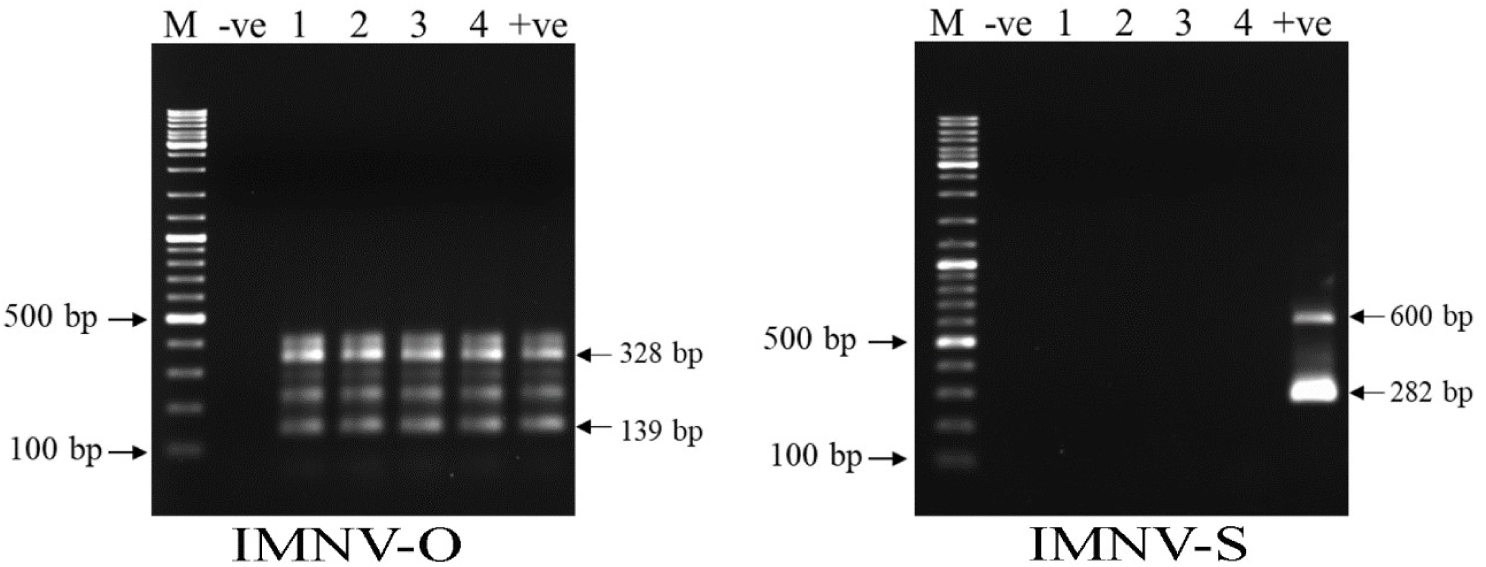
Photographs of agarose gels to detect amplicons from use of the IMNV-O and IMNV-S methods for 4 shrimp specimens (1 to 4) from Lot 4. The 4 specimens were positive with the IMNV-O method but negative with the IMNV-S method. M = molecular marker; -ve = negative control; +ve = positive control.

The IMNV RT-PCR test results with the 4 shrimp in these samples suggested that they do not carry the full genome sequence of what is currently recognized as infectious IMNV. These samples were not studied further. There are several possibilities that may have given rise to these results. The most obvious possibility is that the shrimp may have carried a mutant form of IMNV in which the sequence of the RdRp gene was sufficiently changed to no longer match the sequence of the primers for the IMNV-S method. It is also possible that the positive IMNV-O result arose from an endogenous viral element (EVE) that produced an RNA transcript with sufficient sequence similarity to give amplicons with the method. To check this possibility, we treated DNA extracts from these specimens with RNase and then carried out direct PCR reactions (i.e., no reverse-transcriptase step for cDNA production) with the same IMNV-O primer set, but no amplicons resulted (not shown). We did not sequence the amplicons. However, tissue sections were examined for muscle lesions (see the next section).

### Muscle lesions seen in Lot 4 IMNV PCR-positive samples

Muscle lesions similar to those caused by IMNV were seen in the 4 shrimp samples positive for IMNV by the IMNV-O method. An example is shown in **Fig. 5A**. Most of the lesions did not show hemocytic aggregation. None showed basophilic cytoplasmic inclusions characteristic of IMNV lesions reported for penaeid shrimp including *P. monodon* (Tang, et al., 2005). The problem is that muscle lesions that may or may not cause gross whitening of muscles in living shrimp can be caused simply by stress such as handling (e.g., muscle cramp syndrome or idiopathic myonecrosis) (Lightner, 1988) or by other viruses such as *Macrobrachium rosenbergii* nodavirus (MrNV) (Sahul Hameed, et al., 2004; Sri Widada, et al., 2003) and *Penaeus vannamei* nodavirus (PvNV) (Tang, et al., 2007; Tang, et al., 2011). Thus, further testing by ISH and immunohistochemistry were needed to conform whether the lesions exemplified in Fig. 4A arose from IMNV or not. To this end, tissue sections showing these lesions were tested for the presence of IMNV by ISH using probes for both the IMNV-O target and the IMNV-S target (**Fig. 5C&D**), but all gave negative test results, in contrast to the positive control obtained from Arizona.

**Figure 5.**
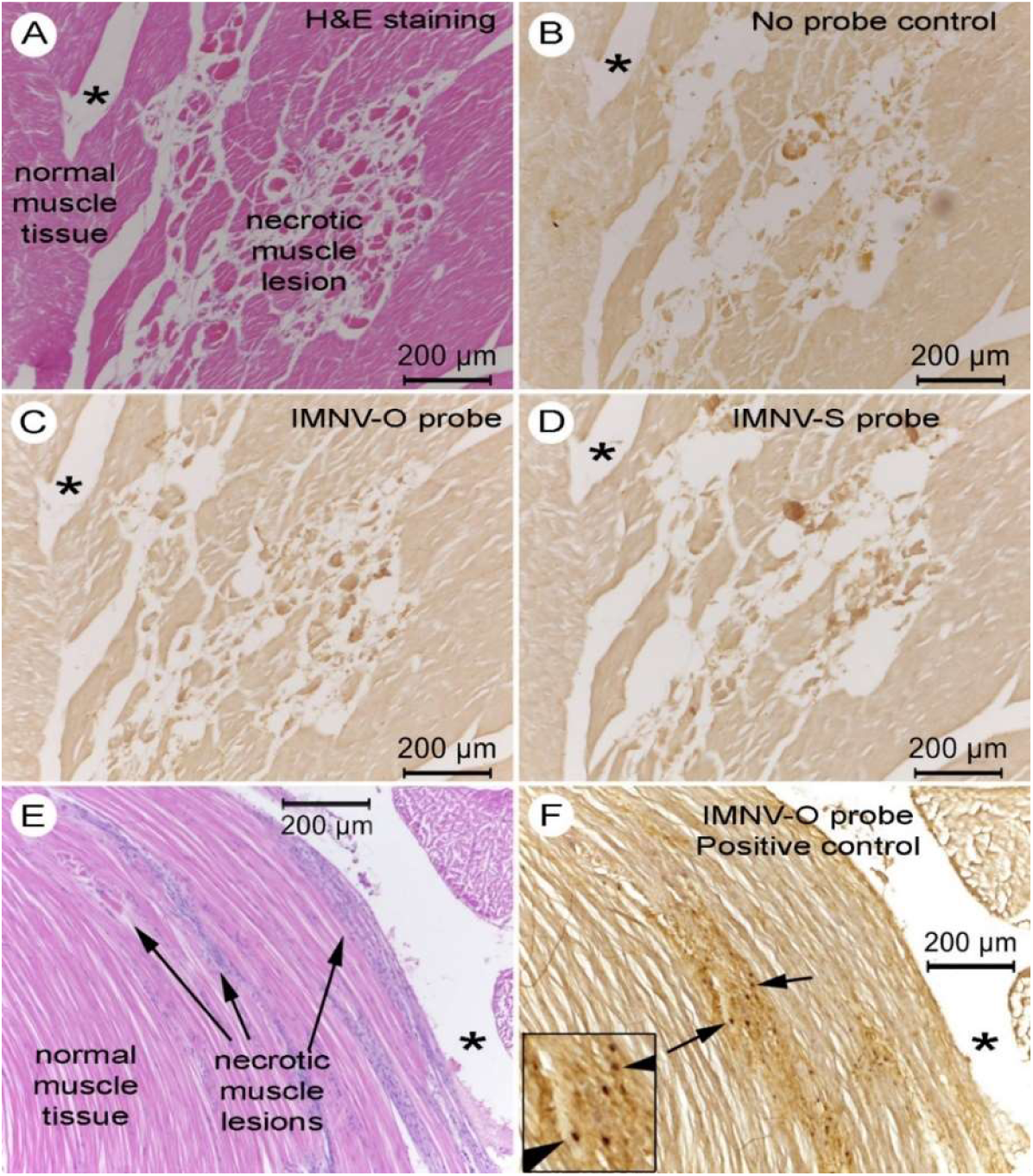
Photomicrographs showing examples of muscle lesions in *P. monodon* samples from Lot 4 positive for IMNV using the IMNV-O method only. Asterisks indicate the same relative position in adjacent tissue sections. (A) H&E stained tissue section showing muscle necrosis. (B) ISH results for the negative (no probe) control showing no signal. (C) ISH result for the IMNV-O probe showing no signal. (D). ISH results for the IMNV-S probe showing no signal. (D) H&E stained tissue section of the positive control tissue showing IMNV muscle lesions. (E) Positive control tissue section showing a positive ISH reaction for IMNV using the IMNV-O method. The inset shows a magnification of area with positive ISH reactions.

### Brief summary for IMNV results

In summary, our results have revealed by 2 RT-PCR methods that some captured specimens of *P. monodon* from the Indian Ocean may carry infectious IMNV. This cannot be confirmed without full genome sequencing and bioassays. However, the results were unexpected and warrant caution in the use of captured broodstock in shrimp hatcheries, and especially those hatcheries that also process *P. vannamei* or produce *P. monodon* PL that are destined for use on farms were *P. vannamei* is also cultivated.

### Positive PCR test results for DIV1

The positive PCR test results for ATPase method were found in both shrimp sample Lot 1 and Lot 4. Shrimp positive ATPase methods were 5 (out of 14) and 8 (out of 76) for Lot 1 and Lot 4, whereas only 1 shrimp from each Lot found positive for both ATPase and MCP (#3 for Lot 1 and #2 for Lot 4) (**Fig. 6**).

**Figure 6.**
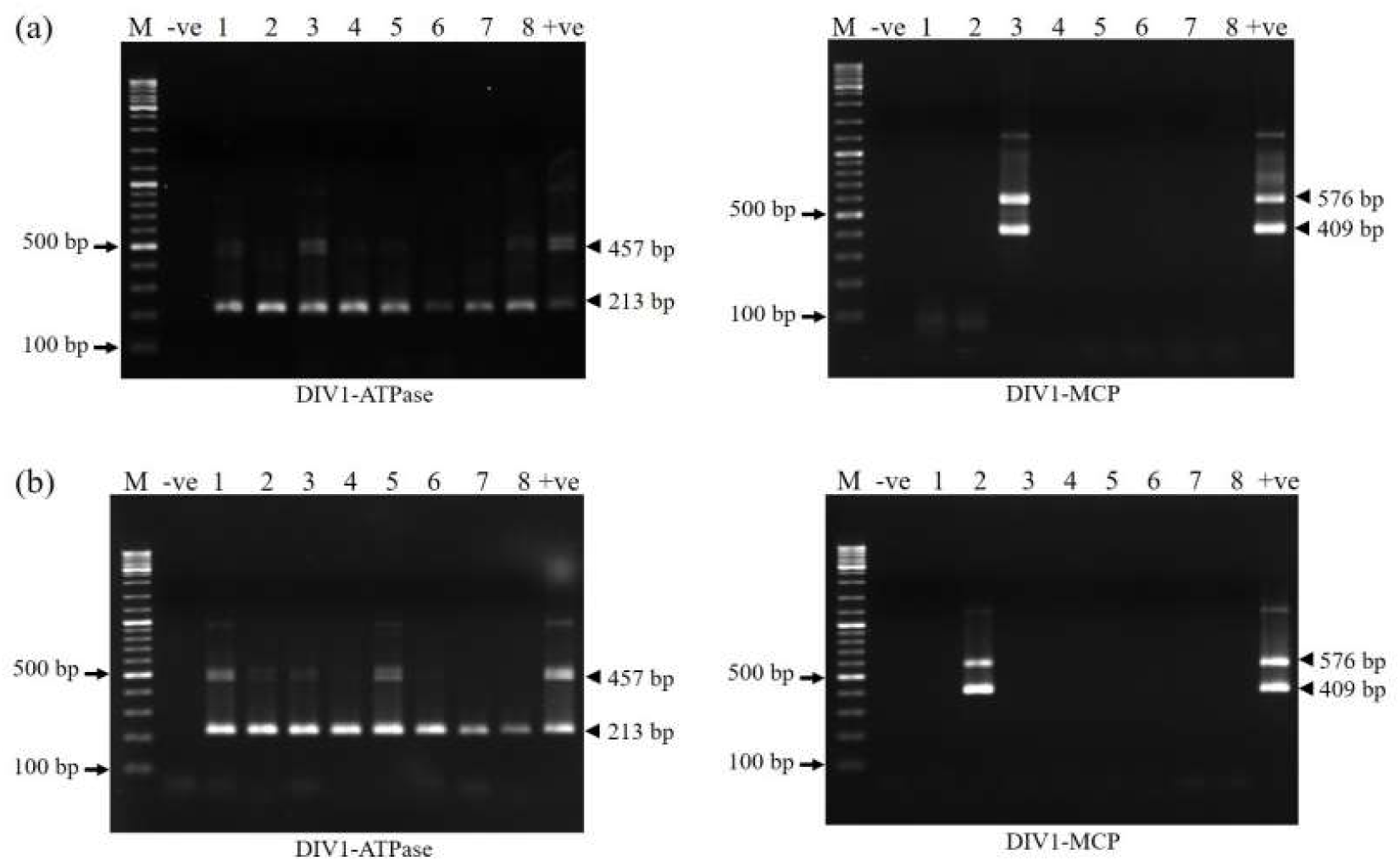
Photograph of the agarose gels showing PCR results for the ATPase and MCP methods in 5 out of 14 in specimen Lot 1 (a) and 8 out of 76 shrimp samples in specimen Lot 4 (b). Only 1 out of 5 specimens with ATPase positive in Lot 1 and 1 out of 8 specimens with ATPase positive in Lot 4 also gave a positive test result with the MCP method.

The PCR results for the ATPase and MCP methods for detection of DIV1 were both positive for only one specimen of each Lot. This suggested that this specimen might have carried DIV1. Thus, the specimens from Lot 4 were subjected to sequencing and analysis of the amplicons from the two PCR methods. As can be seen in **Figs. 7 and 8**, the sequence identities were 100% for both amplicons when compared to the matching regions of the full genome of DIV1 (GenBank KY681040.1).

**Figure 7.**
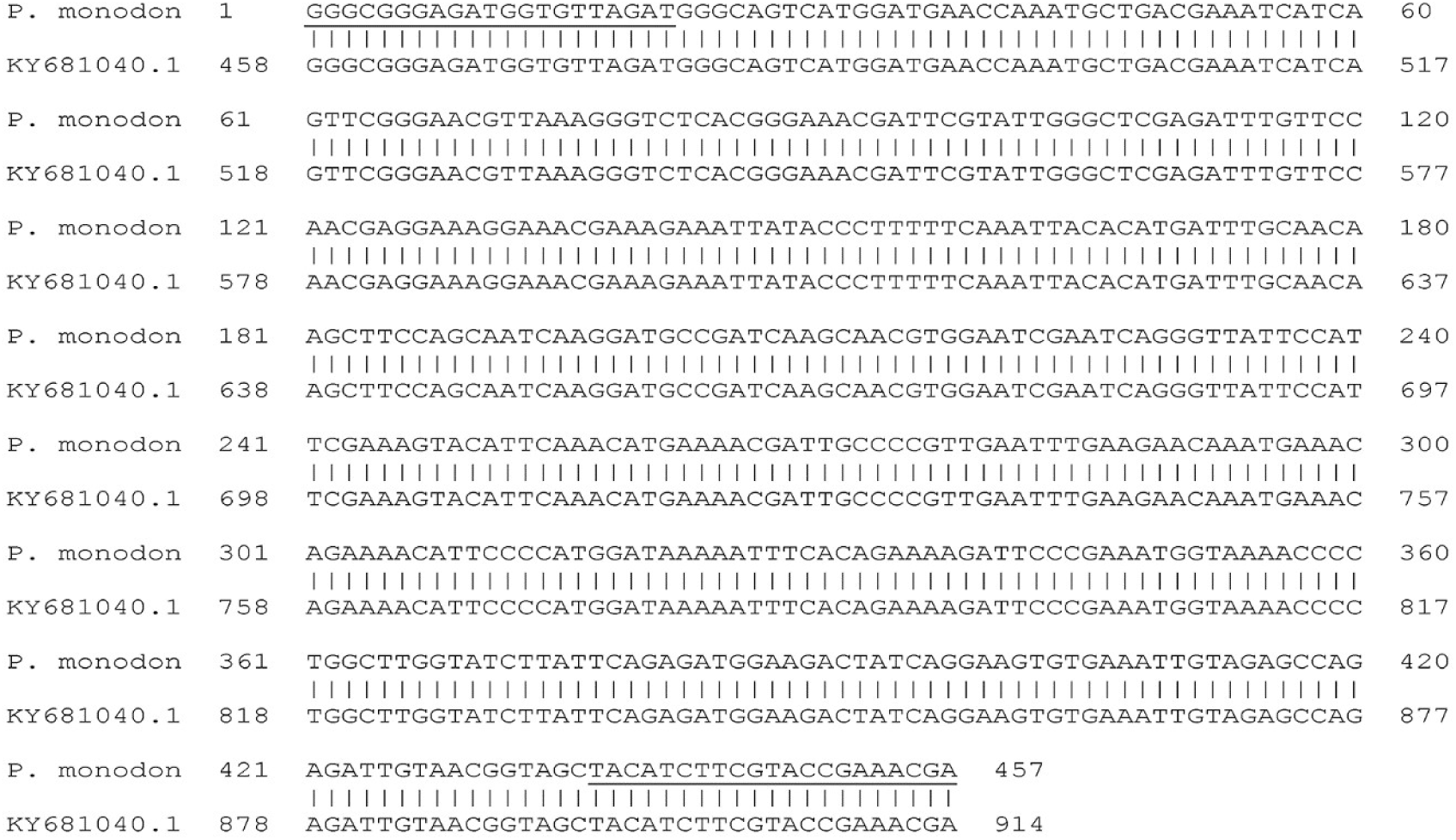
Alignment of the amplicon sequence obtained using the ATPase method compared to the matching region of the full genome of DIV1 (GenBank KY681040.1). The primer sequences are underlined.

**Figure 8.**
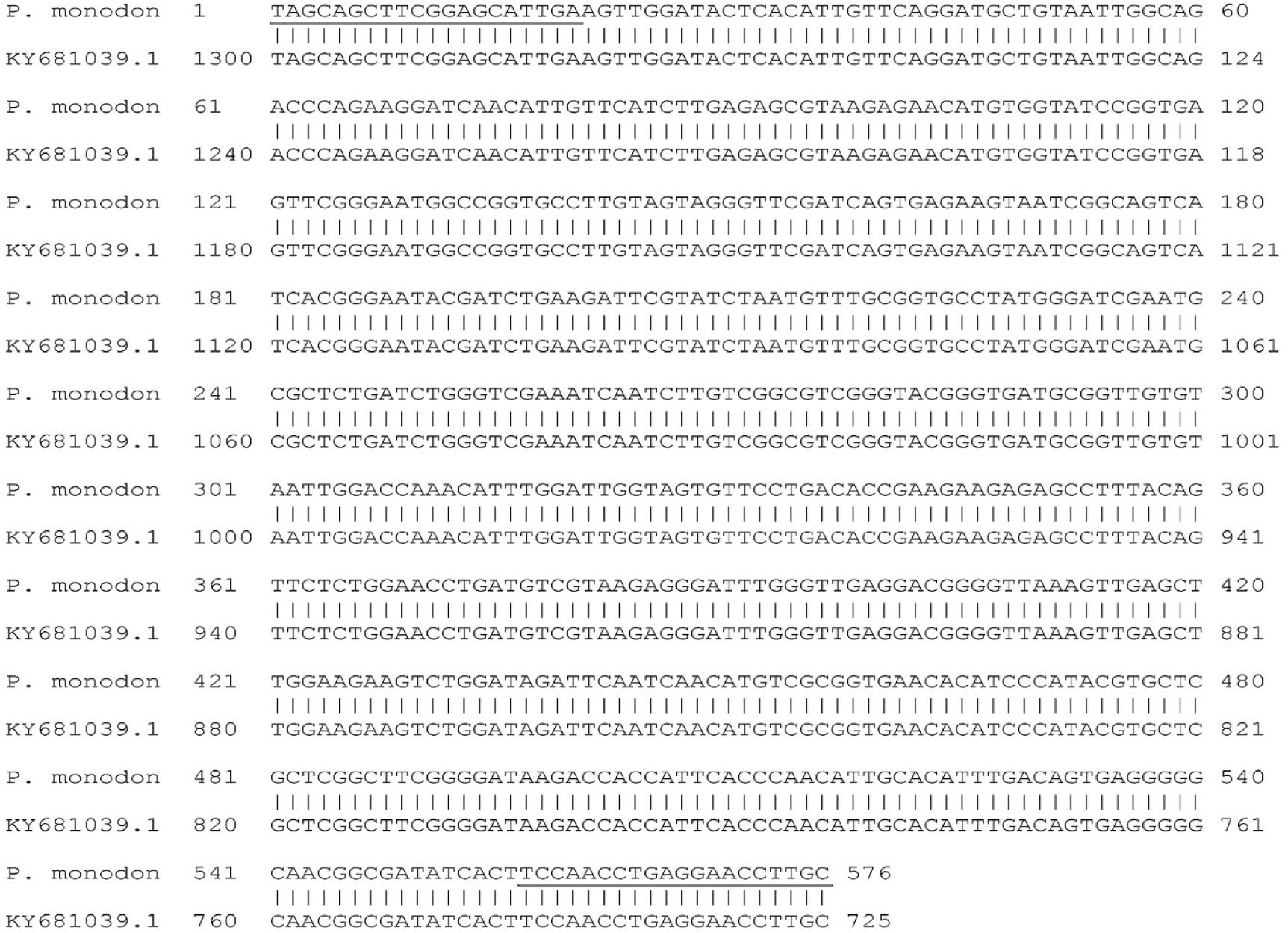
Alignment of the amplicon sequence obtained using the MCP method compared to the matching region of the full genome of DIV1 (GenBank KY681040.1). The primer sequences are underlined,

The PCR positive results and sequencing results for 2 distantly separated genes in the DIV1 genome for 1 out of 76 specimens in shrimp Lot 4 raised the possibility that captured, broodstock-size specimens of *P. monodon* from the Indian Ocean may be infected with a virulent type of DIV1. However, it is important to understand that this possibility cannot be substantiated without full sequencing of the whole viral genome accompanied with bioassays. As with IMNV above, this was an unexpected result, but of sufficient importance to warrant caution in the use of captured *P. monodon* for broodstock in shrimp hatcheries and especially in those hatcheries that also process *P. vannamei* or produce PL for use on farms were *P. vannamei* is also cultivated.

### Lack of gross and histological signs of DIV1 infection

All 8 of the specimens positive for DIV1 in Lot 4 using the ATPase method were grossly normal and showed no gross signs of DIV1 infection, including the 1 specimen also positive for DIV1 using the MCP method. Of the 6 of 8 specimens arbitrarily selected for ISH testing, (including the one positive for both the ATPase and MCP methods), all showed normal HPT histology (**Fig 9A**) and normal LO histology (**Fig. 10A**) except, sometimes for the presence of spheroids in the latter. Such spheroids have not been reported to be associated with DIV1-infection. Those not familiar with the pathognomonic lesions in of DIV1 in the HPT and its lesions in the LO may download the DIV1 disease card for free from the website of the Network of Aquaculture Centres in Asia Pacific (NACA). All the other specimens gave similar results for HPT and LO histology.

**Figure 9.**
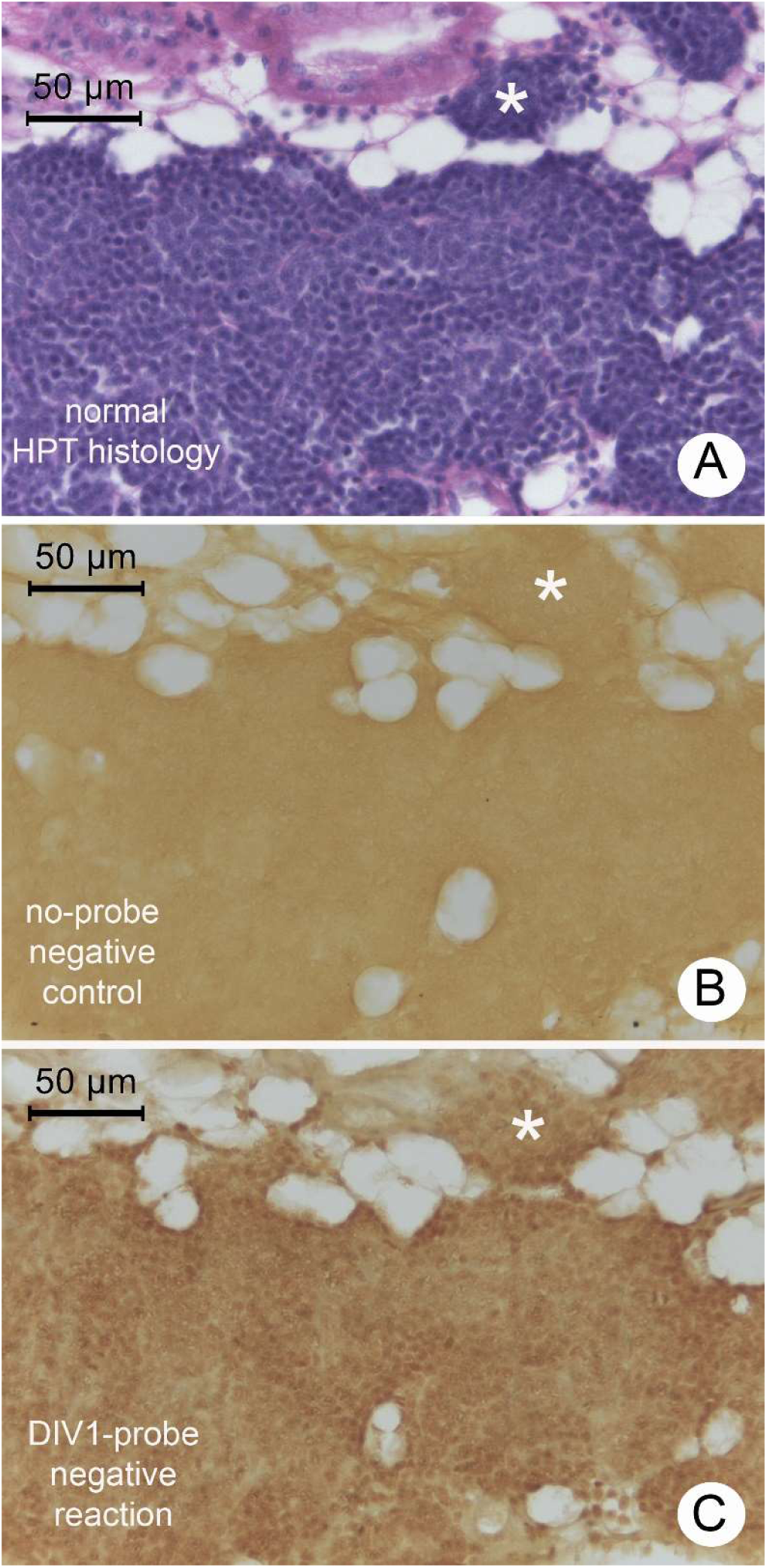
Example photomicrographs of the HPT from one of the 4 PCR-positive DIV1 specimens arbitrarily selected for ISH analysis (including the specimen positive for both target genes). (A) H&E stained HPT section showing normal histology (i.e., absence of DIV1 lesions characterized by lightly basophilic cytoplasmic inclusions). (B) Negative, no-probe control ISH reaction with an adjacent tissue section showing no signals. (C) Negative ISH reaction with the DIV1-DIG-labeled probe for the ATPase gene. Similar results were obtained for all 4 specimens with both the ATPase and the MCP probe.

**Figure 10.**
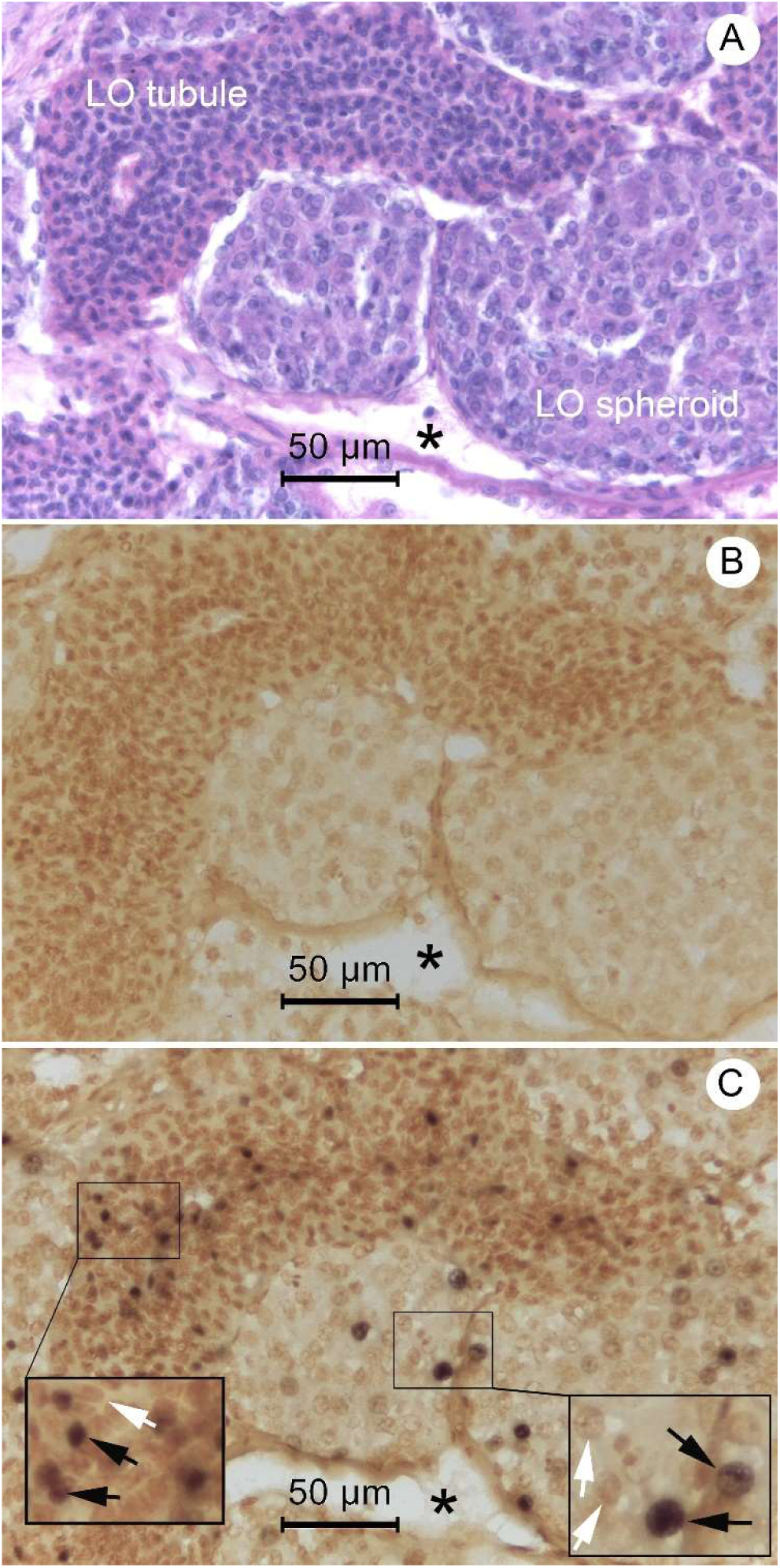
Example photomicrographs the LO tissue one of 4 PCR-positive DIV1 specimens selected for ISH analysis. (A) H&E stained LO section showing normal tubule histology (i.e., absence of DIV1 lesions characterized densely basophilic cytoplasmic inclusions and pyknotic and karyorrhectic nuclei). (B) Negative, no-probe control ISH reaction with an adjacent tissue section showing no signals. (C) Curious ISH reaction with the DIV1-DIG-labeled probe showing dark staining in the nuclei but not the cytoplasm of both tubule matrix cells and cells in spheroids.

Although all the 8 specimens positive for DIV1 by PCR using the ATPase method gave negative RT-PCR test results for IMNV, 5 out of the 6 examined histologically showed necrotic muscle lesions, similar to those seen with the specimens above that gave positive RT-PCR test results but negative ISH test results for IMNV. Thus, the results from the DIV1 positive specimens support the proposal above that the muscle lesions that gave negative ISH results for IMNV were probably expressions of idiopathic myonecrosis (Lightner, 1988).

### Typical, positive ISH results for DIV1 in the LO and HP

Example photomicrographs of unusual ISH test results for DIV1 in the LO and HP tissue of the same 4 shrimp specimens that gave negative ISH reactions for DIV1 in the HPT. These unusual results are shown in **Figs. 10C and 11C** where positive ISH results can be seen in the nuclei of some cells in the LO and HP in the absence of DIV1 histopathology. The same result was obtained in the ISH tests with all 4 samples tested, including 3 positive for DIV1 by PCR with the ATPase method only and 1 positive for DIV1 by PCR with both the ATPase and MCP methods.

An example of the unexpected, positive ISH reactions in many nuclei in both the LO tubule matrix and in the spheroids are shown in **Fig 10C**. In other samples without spheroids, the signals were still present in the nuclei of tubule matrix cells. This contrasts curiously with the situation in DIV1-diseased shrimp where the lesions occur in the LO cell cytoplasm rather than the nuclei and where they are accompanied by densely basophilic, karyorrhectic and pyknotic nuclei that give photomicrographs a “peppered” appearance (see the NACA disease card for DIV1) similar to that seen with lesions of yellow head virus (YHV)(Flegel, 2006).

An example of the unexpected positive ISH reactions in the nuclei of the tubule epithelial cells of the HP of the same 4 specimens as above are shown in **Fig. 11C** Again, the reactions differ from the situation in DIV1-diseased shrimp where the ISH signals arise in the cytoplasm of cells in the interstitial spaces (i.e., connective tissue) that separate the HP tubules. A comparison is shown at high magnification in **Fig. 12**.

**Figure 11.**
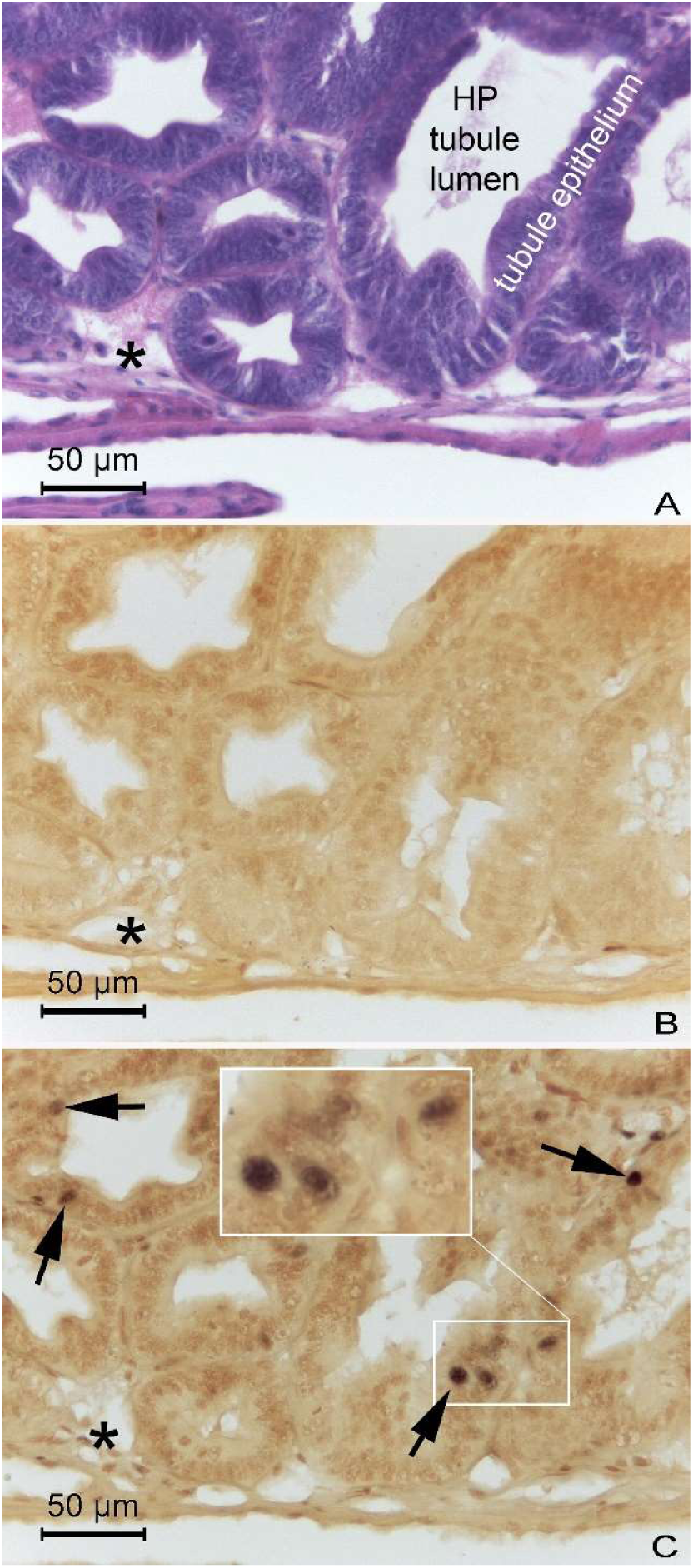
Example of the curious, positive ISH reactions for DIV1 in the nuclei of tubule epithelial cells of the HP. The asterisks in each photomicrograph indicate the same relative position in the 3 adjacent tissue sections. (A) H&E stained section showing normal HP histology. (B) No-probe negative control showing no ISH signals. (C) Positive ISH signals in the nuclei of HP tubule epithelial cells.

**Figure 12.**
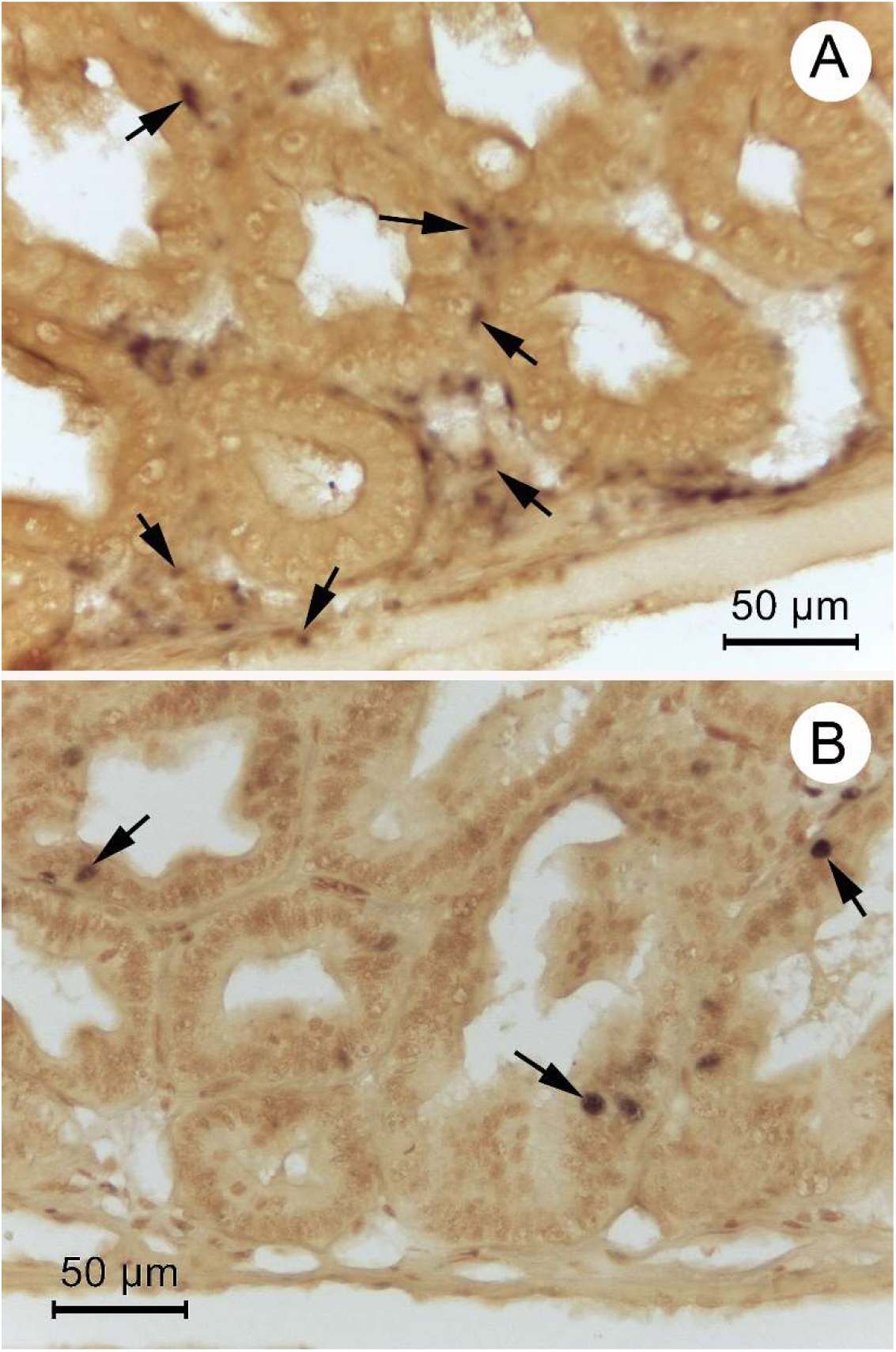
Comparison of photomicrographs showing positive ISH reactions (arrows) for DIV1 in the HP tissue of diseased shrimp (A) and non-diseased shrimp (B), both positive for DIV1 by PCR using the ATPase method. In (A) the positive reactions occur connective tissue in the interstitial spaces between the HP tubules while in (B) (copy from Fig. 10C above) they occur in the nuclei of the tubule epithelial cells.

## DISCUSSION

We used 2 different PCR methods with each virus targeted in this study. The first method for each was the standard one used for each of the viruses in our laboratory. The second method for each virus was not used routinely and was employed in order to further avoid the possibility of contamination from post-PCR material. In addition, the second method for each virus was designed to be distant on the respective genomes from the region used in the first tests, reducing the possibility that amplicons could have arisen from a single incomplete viral segment. The fact that both tests for each virus were positive for the same shrimp specimens in each batch and the fact that the sequence identities were 99-100% for all the target sequences made us confident that the target amplicons did not arise from laboratory contamination. However, we cannot exclude the possibility that the amplified sequences arose from incomplete viral genomes or from endogenous viral elements (EVE) such as have been reported, for example, for infectious hypodermal and hematopoietic necrosis virus (IHHNV) (Saksmerprome, et al., 2011; Tang Lightner, 2006) in *P. monodon* and *P. vannamei* and white spot syndrome virus (WSSV) (Taengchaiyaphum, et al., 2019; Utari, et al., 2017) in *P. monodon*. Thus, full viral genome sequencing and ultimately bioassays would be required to confirm whether the captured *P. monodon* would be capable of transmitting IMNV and DIV1.

The fact that the ISH tests for IMNV were negative weakens the possibility that the specimens were carrying infectious IMNV. However, grossly normal broodstock specimens of *P. vannamei* infected with IMNV and positive for it by nested-RT-PCR often give negative ISH test results but remain able to transmit IMNV to naïve shrimp. On the other hand, 6 out of 8 broodstock specimens in sample Lot 4 that were positive for DIV1 but negative for IMNV using PCR methods showed muscle necrosis similar to those in the specimens positive for IMNV by RT-PCR, indicating that they arose from some other cause. Despite these uncertainties, we believe it is better to be cautious and exclude such shrimp as candidates for PL production.

With respect to the specimens that gave positive PCR results for DIV1, the histological analysis also revealed no pathognomonic lesions in the HPT or supporting lesions in the LO. Nor were there any positive ISH reactions in the HPT, one of its prime target tissues. However, unlike IMNV, some positive but atypical ISH test results were obtained in one of its prime target organs (i.e., in the LO) despite the lack of DIV1-type lesions. They were atypical because the positive signals arose in the nuclei instead of the cytoplasm as is expected for DIV-diseased shrimp. Even more surprising was the occurrence of positive ISH reactions in nuclei of the tubule epithelial cells of the HP. This was surprising not only because the positive signals were from a nucleus but also because positive ISH reactions in the HP of DIV-diseased shrimp occur only in the connective tissue of the HP (i.e., in the interstitial spaces between the tubules). Indeed, there were no positive ISH reactions in other connective tissues of these specimens, even though such reactions are widespread in the connective tissue of DIV-diseased shrimp

The atypical ISH reactions for DIV1 in the captured broodstock specimens when compared to those in DIV-diseased shrimp raise many questions. None of these can be answered without further research. The most important questions for shrimp farmers are those related to the possibility of DIV1 disease transmission. For example, it is known that iridoviruses have both nuclear and cytoplasmic stages in the life cycle and that most of the viral production occurs in the cytoplasm. This corresponds to the experience in DIV1-diseased shrimp where strong ISH reactions have been reported in the cytoplasm where masses of virions can be seen by transmission electron microscopy. However, positive ISH reactions have not been reported in the nuclei of DIV1-diseased shrimp, so it is possible that the amount of viral DNA in the nuclei during the disease state is insufficient to give a visible signal with the methods that have been used. Is it possible that the positive ISH reactions seen in our captured specimens indicate the presence of an inactive stage of DIV1 that has persisted in survivors from a previous exposure? If so, such hosts might serve as tolerant carriers. Yet another possibility is that the specimens we examined carried a non-pathogenic, genetic variant of DIV1 that differed from the type described from China. Again, only complete genome sequencing and bioassays could answer these questions. It is also possible that *P. monodon* is tolerant to DIV1 and carries it in a latent state in nuclei of the HP and LO.

Because of the uncertainties discussed above, we cannot confidently dismiss the possibility that the grossly normal, PCR-positive, captured *P. monodon* specimens we examined might have been infected with the respective viruses at the carrier level. If so, they might serve as potential vehicles for introduction of IMNV and/or DIV1 into crustacean culture systems, especially if they were used in hatcheries for production of PL for distribution to shrimp farmers without proper precautions in place. It is already known that *P. monodon* may be infected with IMNV without showing gross signs of disease (Tang, et al., 2005) and our results suggest that the long presence of IMNV in Indonesia after its introduction around 2007 (Senapin, et al., 2007) may have resulted in its transfer from shrimp farms to grossly normal wild stocks of *P. monodon*. If this is so and if infectious IMNV is present in a significant portion of *P. monodon* in the Indian Ocean, it is possible that the recent outbreak of IMNV at a *P. vannamei* farm in Malaysia in June 2018 (WAHID, OIE) might have occurred via this transmission pathway.

Although the presence of IMNV in wild *P. monodon* may be proposed to have arisen because of its long presence in Indonesia after introduction there around 2006, it is more difficult to hypothesize the pathway for occurrence of DIV1-positive specimens because the virus was first described from China less than 4 years ago (Qiu, et al., 2017; Xu, et al., 2016). If DIV1 is a newly emerging pathogen from China it seems unlikely that it could have spread to the Indian Ocean and reached a significant presence in the wild *P. monodon* population there within 3 years simply by movement of wild, infected shrimp. It also seems unlikely that DIV1 could have been endemic in *P. monodon* but been overlooked or not have caused any mortality in exotic *P. vannamei* since it became the dominant cultivated species from the early 2000’s onward. This is especially so when one considers that *P. vannamei* went through several years of cultivation near and together with *P. monodon* without DIV1 disease outbreaks before and even after *P. vannamei* grew to dominance.

In summary, because we did not amplify and sequence the whole viral genome from the specimens, and because we did not do bioassays, we cannot confirm that the shrimp were carriers of infectious IMNV or DIV1. However, we believe that our PCR results justify a precautionary warning regarding the possibility of introducing IMNV and DIV1 into aquaculture facilities via use of wild, captured *P. monodon* from the Indian Ocean. To avoid this possibility, we recommend that wild, captured *P. monodon* from the Indian Ocean intended for use as broodstock be subjected to PCR testing for DIV1 and IMNV before use in a hatchery and that they be discarded, if they are found to be positive. If not positive, their larvae and post-larvae (PL) should be monitored for presence of these 2 viruses periodically during production and again before they are sold to users. We also strongly recommend that industry practitioners who currently use wild, captured *P. monodon* be discouraged from handling them together with broodstock of other crustaceans listed above in common maturation or hatchery facilities. In addition, we recommend that shrimp farmers be discouraged from cultivating those species together with *P. monodon* in the same pond or on the same farm, especially if the latter originated from wild, captured broodstock that have not been tested for freedom from IMNV and/or DIV1, as applicable based on susceptibility of the specific species. Indeed, since domesticated stocks of *P. monodon* SPF for IMNV and DIV1 are available, we do not recommend the use of captured wild *P. monodon* broodstock for PL production at all. One reason is to prevent not only transmission of these two viruses, but also monodon baculovirus (MBV), hepatopancreatic parvovirus (HPV), white spot syndrome virus (WSSV) and yellow head virus (YHV). In addition, reducing the fishing pressure on shrimp broodstock should help to promote a more sustainable natural shrimp fishery.

## Acknowledgement

We would like to thank the financial support from the Newton prize (Chairman’s award) 2017 and the Royal Society International Collaboration Awards 2019, the Global Challenges Research Fund (GCRF) to Prof. Grant D. Stentiford, Cefas/UK and Dr. Kallaya Sritunyalucksana, BIOTEC, NSTDA/Thailand.

